# Development of a SARS-CoV-2 monoclonal antibody panel and its applicability as a reagent in high-throughput fluorescence reduction neutralization and immunohistochemistry assays

**DOI:** 10.1101/2023.05.29.542720

**Authors:** Gabriela Mattoso Coelho, Allan Henrique Depieri Cataneo, Sonia Mara Raboni, Meri Bordignon Nogueira, Caroline Busatta Vaz de Paula, Ana Clara Simões Flórido Almeida, Vanessa Zulkievicz Rogerio, Nilson T. Zanchin, Lucia de Noronha, Camila Zanluca, Claudia Nunes Duarte dos Santos

## Abstract

Since its emergence in late 2019, infection by SARS-CoV-2 (COVID-19 disease) has quickly spread worldwide, leading to a pandemic that has caused millions of deaths and huge socio-economic losses. Although vaccination against COVID-19 has significantly reduced disease mortality, it has been shown that protection wanes over time, and that circulating SARS-CoV-2 variants may escape vaccine-derived immunity. Therefore, serological studies are still necessary to assess protection in the population and better guide vaccine booster programs. A common measure of protective immunity is the presence of neutralizing antibodies (nAbs). However, the gold standard method for measuring nAbs (plaque reduction neutralization test, or PRNT) is laborious and time-consuming, limiting its large-scale applicability. In this study, we developed a high-throughput fluorescence reduction neutralization assay (FRNA) to detect SARS-CoV-2 nAbs. Because the assay relies on immunostaining, we also developed and characterized in-house monoclonal antibodies (mAbs) to lower assay costs and reduce the vulnerability of the test to reagent shortages. Using samples collected before the pandemic and from individuals vaccinated against COVID-19, we showed that the results of the FRNA we developed using commercial and in-house mAbs strongly correlated with those of the standard PRNT method while providing results in 70% less time. In addition to providing a fast, reliable, and high-throughput alternative for measuring nAbs, the FRNA can be easily customized to assess other SARS-CoV-2 variants of concern (VOCs).

## Introduction

Over the last three years, the world has faced the coronavirus disease 2019 (COVID-19) pandemic caused by the newly identified beta-coronavirus SARS-CoV-2^1^. Since its emergence in late 2019 in Wuhan, China, the virus has quickly spread worldwide through interhuman transmission, causing the collapse of healthcare systems in many countries and extensive socio-economic losses. As of December 2022, the pandemic has affected 230 countries and territories, with more than 680 million confirmed infections and over 6.8 million deaths^2^. In addition, China has recently faced a rapid surge of COVID-19 cases following the end of its zero-Covid policy in late 2022.

Infection by SARS-CoV-2 was initially described as being a respiratory disease that commonly causes cough, fever, fatigue, and shortness of breath. A progression to serious complications such as severe pneumonia or acute respiratory distress syndrome (ARDS) was more likely to develop in patients with comorbidities such as diabetes, obesity, heart conditions, and chronic lung disease^3,4^. As the pandemic proceeded, it was shown that COVID-19 affects other organ systems, including the cardiovascular, gastrointestinal, and neurological systems and is therefore a systemic disease. Moreover, many recovered patients deal with long-term symptoms and sequelae after acute infection, known as “long COVID”^5,6^.

Fortunately, global efforts have led to the development of several COVID-19 vaccines, which contributed significantly to reducing viral transmission, hospitalizations, and mortality^7^. Even with the availability of licensed vaccines of proven efficacy, follow-up serological studies are still necessary to assess the immunological status of the population. The duration of immune protection in vaccinated and COVID-19-recovered individuals is still poorly understood, and a significant portion of the world’s population remains unvaccinated due to uneven vaccine distribution and growing anti-vaccine movements. In addition, the emergence of SARS-CoV-2 variants of concern (VOCs) that may escape the protection offered by current vaccines highlights the need to optimize vaccination strategies through booster shots or the development of updated, customized vaccines^8,9^.

A common assessment of protective immunity is the presence of neutralizing antibodies (nAbs) in the individual’s blood, which are elicited through natural infection or vaccination and can block viral entry in host cells^10^. NAbs are key players in conferring protection against SARS-CoV-2 reinfections and/or severe outcomes and have been proposed as predictors of protection against SARS-CoV-2 variants^11^. Given the correlation of nAbs with immune protection, there is an urgent need for assays that rapidly detect SARS-CoV-2 nAbs which can be used in large-scale seroprevalence studies and vaccine trials. The golden standard method for detecting nAbs is the plaque reduction neutralization test (PRNT). However, it is labor-intensive, time-consuming, and requires large sample volumes, limiting its use in large-scale settings.

In this study, we describe a high-throughput FRNA for detecting SARS-CoV-2 nAbs. This assay overcomes the limitations of a classical PRNT assay, allowing for faster results and scalable testing of large numbers of samples simultaneously. To reduce assay costs, we develop anti-SARS-CoV-2 mAbs and demonstrate their applicability in the FRNA and immunohistochemistry assays, comparing their performance with commercially available mAbs.

## Methods

### Cell lines and viruses

Vero E6 cells from *Cercopithecus aethiops* (85020206; Sigma, St. Louis, MO, USA) and Vero E6-TMPRSS2 (kindly provided by Dr. José Luis Proença Módena) were cultivated in Dulbecco’s Modified Eagle Medium: Nutrient Mixture F-12 (DMEM F-12) supplemented with 10% heat-inactivated fetal bovine serum (FBS), 100 IU/mL penicillin and 100 μg/mL streptomycin (all reagents from Gibco, Grand Island, NY, USA). Myeloma cell line P3X63Ag8.653 from *Mus musculus* (ATCC CRL-1580) and hybridomas were cultivated in RPMI-1640 medium (Gibco, Grand Island, NY USA) with 20% FBS, 9.6 mM HEPES, 2 mM L-glutamine, 1 mM sodium pyruvate, 100 IU/mL penicillin, 100 μg/mL streptomycin and 0.25 µg/mL amphotericin B. All cells were kept at 37°C in a humidified, 5% CO_2_-controlled atmosphere and were passaged twice a week. The SARS-CoV-2 isolates used in the study were obtained from clinical specimens (nasopharyngeal swabs) of patients with laboratory diagnoses of COVID-19 (positive RT-PCR tests). For virus isolation, Vero E6 cells were previously seeded in 25 cm^2^ flasks, and 300 μL of the clinical sample was filtered in 0.22 μm pore filters before being incubated with the cells at 37°C with 5% CO_2_. After 1h, DMEM-F12 medium was added, and cells were monitored daily for cytopathic effect (CPE). Once CPE was observed, virus isolation was confirmed by RT-PCR and later by NGS sequencing. To prepare viral stocks, Vero E6 cells were infected with a multiplicity of infection (MOI) of 0.001 and harvested after 72h. Stocks were titrated by plaque assay in Vero E6 cells.

### Human serum samples

The study was approved by Fiocruz and the Brazilian National Ethics Committee for Human Experimentation (CAAE 34734920.6.0000.5248). Human serum samples from 50 individuals vaccinated against COVID-19 with the BNT162b2 (Pfizer-BioNTech), AZD1222 (Oxford-AstraZeneca), and CoronaVac (Sinovac Biotech) vaccines were collected at three different time points: T0 (before vaccination), T1 (15 days after the first vaccine dose) and T2 (15 days after the second dose). A panel of 24 serum samples collected before the COVID-19 pandemic was also tested to establish a cutoff value for the assay. All human sera were heat inactivated at 56°C for 30 min before being used in the neutralization assays.

### Fluorescence reduction neutralization assay

Serum samples were serially diluted between 1:20 and 1:2,560 in DMEM F-12 with 100 IU/mL penicillin and 100 µg/mL streptomycin. Each dilution was mixed with 450 PFU of SARS-CoV-2 stock and incubated at 37°C for an hour. The serum and virus mixtures were then added to Vero E6 cells previously seeded in 96-well plates at a density of 5×10^4^ cells/well and were incubated at 37°C and 5% CO_2_. After one hour, the inoculum was removed, and cells were maintained at 37ºC and 5% CO_2_ for 24h.

Cells were then fixed with methanol/acetone (1:1 v/v) for at least one hour at - 20°C, and infection was detected with an indirect immunofluorescence assay (IFA). Briefly, plates were incubated at 37°C for 30 minutes in blocking buffer (PBS with 1% BSA, 5% FBS, and 1% human AB serum (Lonza, Walkersville, MD, USA), washed three times in PBS with 0.05% tween 20 and incubated with a commercial anti-SARS-CoV-2 Spike antibody (1/1000) (MP Biomedicals, Irvine, CA, USA) or the in-house anti-SARS-CoV-2 mAb (1/400). After 40 minutes of incubation at 37°C, wells were rewashed and incubated with an anti-mouse IgG Alexa Fluor 488 conjugated antibody (1/800) (Thermo Fisher Scientific, Grand Island, NY, USA) and DAPI (4,6-diamidino-2-phenylindole; Invitrogen, Waltham, MA, USA) for nuclei staining. IFA images were captured by an Operetta CLS High-Content Imaging System (Perkin Elmer, Waltham, MA, USA) with a 20x objective. Images were then analyzed with the Harmony High-Content Imaging and Analysis Software (Perkin Elmer, Waltham, MA, USA) to calculate the percentage of infected cells for each plate well.

The Z’ factor (Z’ = 1– [3(σp+σn)/(μp-μn)], where σ is the standard deviation and μ is the mean of positive (p) and negative (n) controls) was calculated for the assessment of assay quality^12^. Results lower than 0.5 were not considered for calculating neutralizing titers. Using the software Prism (GraphPad), infection was normalized by positive and mock controls, and neutralization curves were obtained with a log(inhibitor) vs. normalized response—variable slope model to calculate anti-SARS-CoV-2 neutralization antibody titers (NT_50_), defined as the serum dilution that inhibited 50% of viral infection.

### Comparative performance of FRNA and PRNT

PRNT was used to compare the performance of the FRNA we developed. To this end, serum samples were diluted as previously described, mixed with 150 PFU of SARS-CoV-2 stock, and incubated at 37°C for an hour. The serum and virus mixtures were then added to Vero E6 cells previously seeded in 24-well plates at a density of 3×10^5^ cells/well and were incubated at 37°C with 5% CO_2_. After an hour, the inoculum was removed and replaced with an overlay (1.6% carboxymethyl cellulose and 2% FBS in DMEM/F-12 medium). Cells were maintained at 37°C with 5% CO_2_ for six days and later fixed with 3% paraformaldehyde. Wells were stained with 0.75% crystal violet to visualize plaques, and antibody titer (PRNT50) was determined as the serum dilution that inhibited 50% plaque formation compared to the positive control (virus inoculum without serum).

### Production and characterization of anti-SARS-CoV-2 antibodies

Three young adult (30-45 days) Balb/C mice were immunized to obtain hybridomas secreting anti-SARS-CoV-2 antibodies. The virus used for immunization was obtained from the supernatant of Vero E6 infected cells, which was concentrated through PEG8000 precipitation, purified by sedimentation through a 30%/60% sucrose cushion, and inactivated by ultraviolet light irradiation. The immunization protocol consisted of five doses of 8.3×10^4^ PFU/dose/animal of SARS-CoV-2, intraperitoneally (doses 1-4, with Alu-Gel-S adjuvant) and intravenously (dose 5, without adjuvant), with two-week intervals between each dose. All animal procedures were approved by the Ethical Committee on Animal Research of Fiocruz under protocol LW-27/19. Balb/C mice were maintained at the Animal Facility of the Instituto Carlos Chagas/Fiocruz-PR during immunization with *ad libitum* feeding and a 12h light/dark cycle.

Mice were euthanized three days after the last dose, and their splenocytes were fused with myeloma P3X63Ag8.653 cells, as previously described^13^. Hybridoma selection was carried out for 14 days by adding HAT (100 mM hypoxanthine, 0.4 mM aminopterin, and 16 mM thymidine; Sigma, St. Louis, MO, USA) to RPMI-1640 medium 24 hours after cell fusion. The culture medium was then replaced with RPMI-1640 containing HT (hypoxanthine and thymidine; Sigma, St. Louis, MO, USA), and hybridomas were screened by IFA to detect the production of anti-SARS-CoV-2 antibodies. Positive hybridomas were cloned by limiting dilution to obtain antibodies of monoclonal origin (mAbs). MAbs were then concentrated through ammonium sulfate precipitation, and mAb isotypes were determined using SBA Clonotyping System-HRP (Southern Biotech, Birmingham, AL, USA), according to the manufacturer’s protocol.

The specificity of the mAbs to viral proteins was determined by western blot (WB), using a lysate of VERO E6 cells infected with SARS-CoV-2 and the following recombinant proteins: spike S1 subunit, envelope, nucleocapsid (all from Thermo Fisher Scientific, Grand Island, NY, USA) and the receptor binding domain of SARS-CoV-2 (RBD2) and of SARS-CoV (RBD1). The receptor binding domain of SARS-CoV-2 and of SARS-CoV were produced in house. Synthetic genes encoding the respective RBDs (residues 319-533 of SARS-CoV-2 spike protein of the Wuhan strain, accession number QHD43416.1 and, residues 318-513of SARS-CoV spike protein, accession number AAP33697.1), were acquired from GenScript (Piscataway, NJ, USA) and subcloned into the *Nhe*I and *Xho*I restriction sites of plasmid pIRES2-EGFP (Clontech, Takara Bio USA, Mountain View, CA, USA). The coding sequences of the RBDs contain the secretion signal of the chicken receptor-type tyrosine phosphatase μ (RPTPμ) at the N-terminal and a deca-histidine tag preceded by a TEV protease recognition site in the C-terminal. Plasmids pIRES2-EGFP carrying RBD2 and RBD1 were transfected into HEK293 cells. After selecting stable cell lines, the cultures were expanded and both RBD2 and RBD1 were purified from the culture medium by standard immobilized metal chromatography on a HisTrap FF Crude 5 mL column using an ÄKTA Pure M25 chromatography system (Cytiva, Marlborough, MA, USA).

Viral proteins and cell lysates were loaded into 13% SDS PAGE gels and transferred to PVDF (Polyvinylidene fluoride) membranes (Cytiva, Marlborough, MA, USA). Membranes were incubated in blocking buffer (5% non-fat milk, 20 mM Tris, 137 mM NaCl, pH 7.6), followed by concentrated mAb supernatant as the primary antibody. An anti-histidine tag antibody (Sigma, St. Louis, MO, USA) and serum from the immunized mice were used as positive controls. Additionally, a pre-immune serum and anti-zika mAb were used as negative controls. Membranes were then incubated with an alkaline phosphatase-conjugated anti-mouse IgG (Promega, Madison, WI, USA) as the secondary antibody, and the reaction was revealed with a solution of NBT (nitroblue tetrazolium) and BCIP (5-bromo-4-chloro-3-indolyl-phosphate) (Promega, Madison, WI, USA).

### Immunohistochemistry assay (IHC)

Twenty-four post-mortem lung samples from patients who died from COVID-19 between April and August 2020 were collected for the immunohistochemistry assay. The patients were tested for SARS-CoV-2 infection using nasopharyngeal swabs taken during their ICU hospitalization. RT-qPCR was performed on all patients using the SuperScript−III Platinum® One-Step qRT-PCR Kit (Invitrogen, Waltham, MA, USA). The assay was preceded by making multi-sample paraffin tissue blocks (TMA or Tissue Microarray). The immunohistochemistry technique was performed with a commercial antibody anti-SARS-CoV-2 spike protein (Abcam, Cambridge, UK) and the in-house mAb 1F7. The primary antibodies were incubated overnight as recommended for the technique, and the secondary polymer (Mouse/Rabbit PolyDetector DAB HRP Brown, BSB0205, BioSB, Santa Barbara, CA, USA) was added to the material at room temperature. The reaction was revealed by adding the 2,3-diamino-benzidine complex + hydrogen peroxide substrate. Positive and negative controls were used to validate the reactions, and the slides were scanned using the Axio Scan.Z1 Scanner (ZEISS, Jena, Germany).

## Results

### Development of the FRNA

Figure 1 summarizes the FRNA workflow. Several parameters were initially defined to develop the assay, including the cell line and cell density, time of infection, and MOI. Vero E6 cells were chosen for the assay due to their susceptibility to isolation and propagation of SARS-CoV-like viruses and to support viral replication to high titers in short periods of time^14^. To avoid biased results, the wells of the outer rows and columns of the FRNA plates were not used due to the “edge effect,” which results in heterogeneous cell growth and may impact infection frequencies. Instead, cells were seeded only on the inner wells of the plates, and only culture medium was added to the edges.

**Figure 1.**
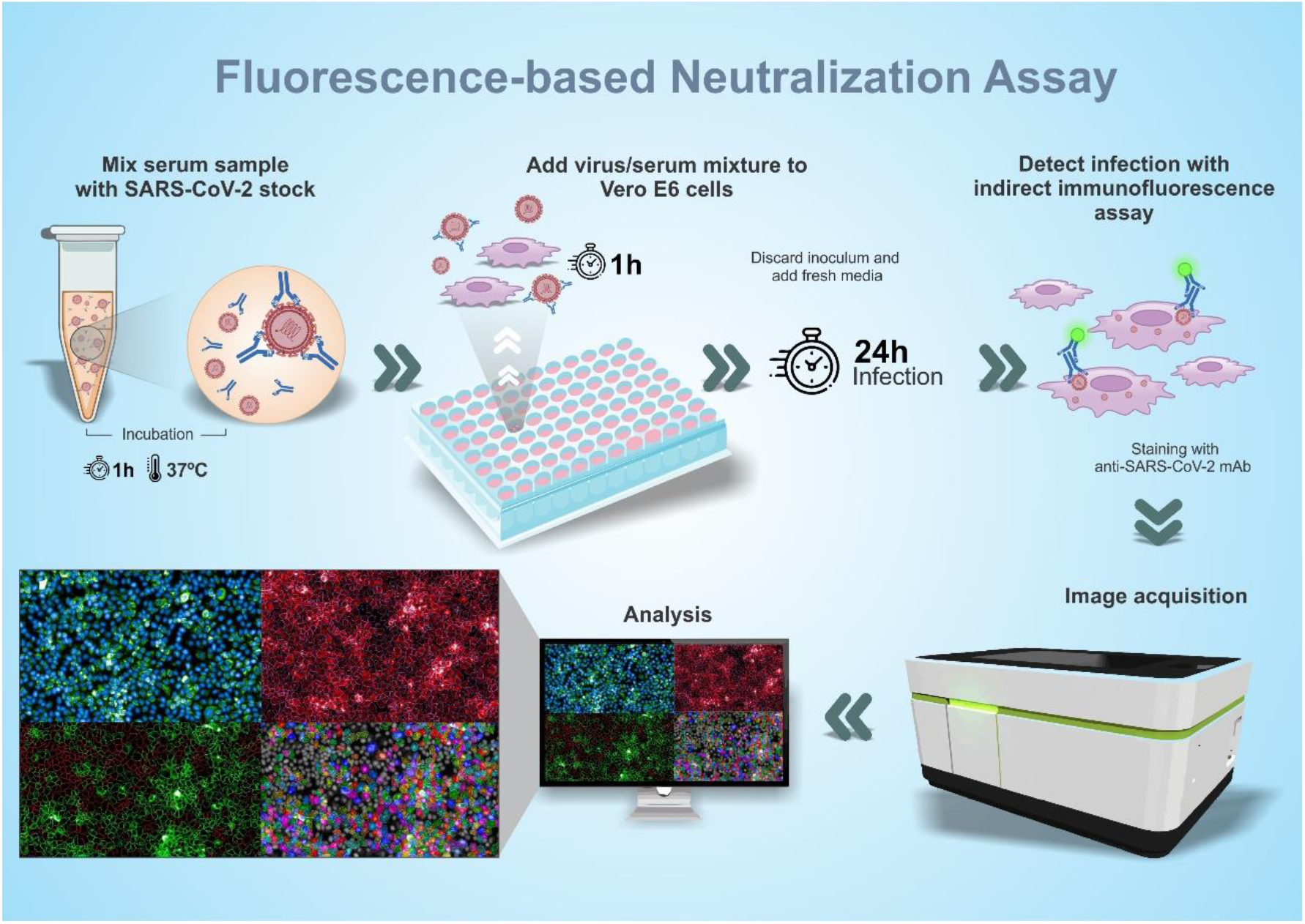
Schematic design illustrating the general procedures used in the assay.

In order to determine the infection parameters, Vero E6 cells were incubated with 450 plaque-forming units of SARS-CoV-2 (MOI 0.009) and analyzed at 20, 24, and 30 hours post-infection (hpi). Following analysis with imaging software, 24hpi was defined as an appropriate time point for subsequent experiments because it resulted in approximately 70% of infected cells with no significant CPE and high Z’ value (Figure 2A). Once we defined the infection parameters, a pre-pandemic and a post-vaccinated sample were used to standardize and calculate the neutralization titers based on a curve-fitting model (Figure 2B and 2C). The curve-fitting model and images obtained from neutralization standardization demonstrated that all parameters set enable further validation.

**Figure 2.**
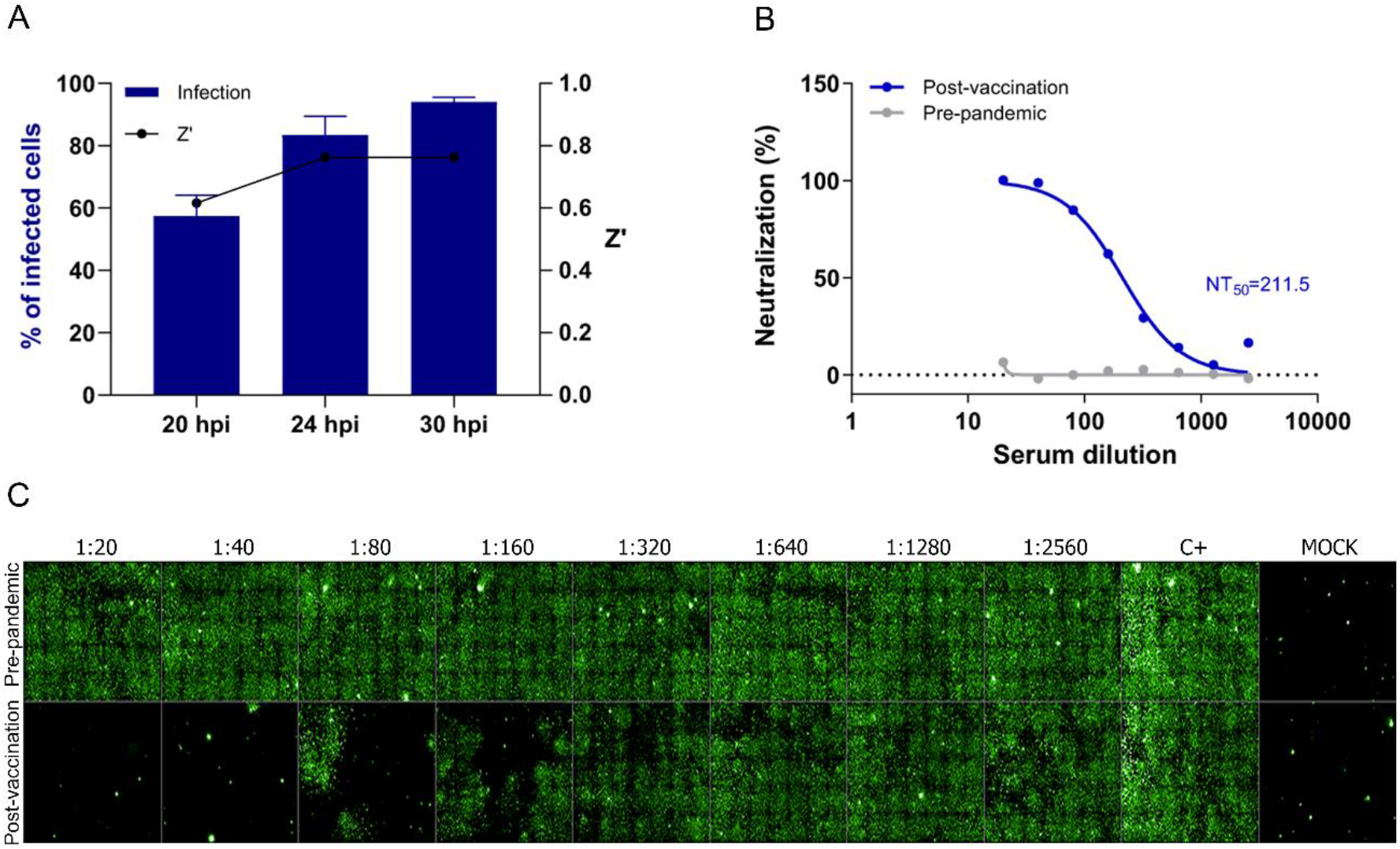
Overview of the standardization of the FRNA for SARS-CoV-2. A) Analysis of the frequency of infected cells and Z’ value at different time points. B) Curve fitting model of results and calculation of neutralization titer inhibiting 50% of SARS-CoV-2 infection (NT50). C) Representative image of serial dilution (1:20 to 1:2560) of pre-pandemic and post-vaccinated samples.

### FRNA validation

The assay validation was performed using a panel of 50 serum samples collected from individuals who received two doses of COVID-19 vaccines BNT162b2, AZD1222, or CoronaVac (i.e., collected at T2). Additionally, 24 pre-pandemic serum samples were tested to establish a cutoff value for the assay. All analyses were performed after checking pre-defined criteria, such as no CPE, appropriate infection rate, and Z’ index above 0.5.

Based on the titers obtained from pre-pandemic serum specimens, samples were considered positive if NT50 was higher than 20, which was the lowest serum dilution used in the assay. This cutoff resulted in a positive neutralization rate of 88% of the serum samples from vaccinated individuals. Antibody titers varied greatly across positive samples, with NT50 ranging from 25 (lowest titer) to 10,000 (highest titer obtained) (Figure 3A).

**Figure 3.**
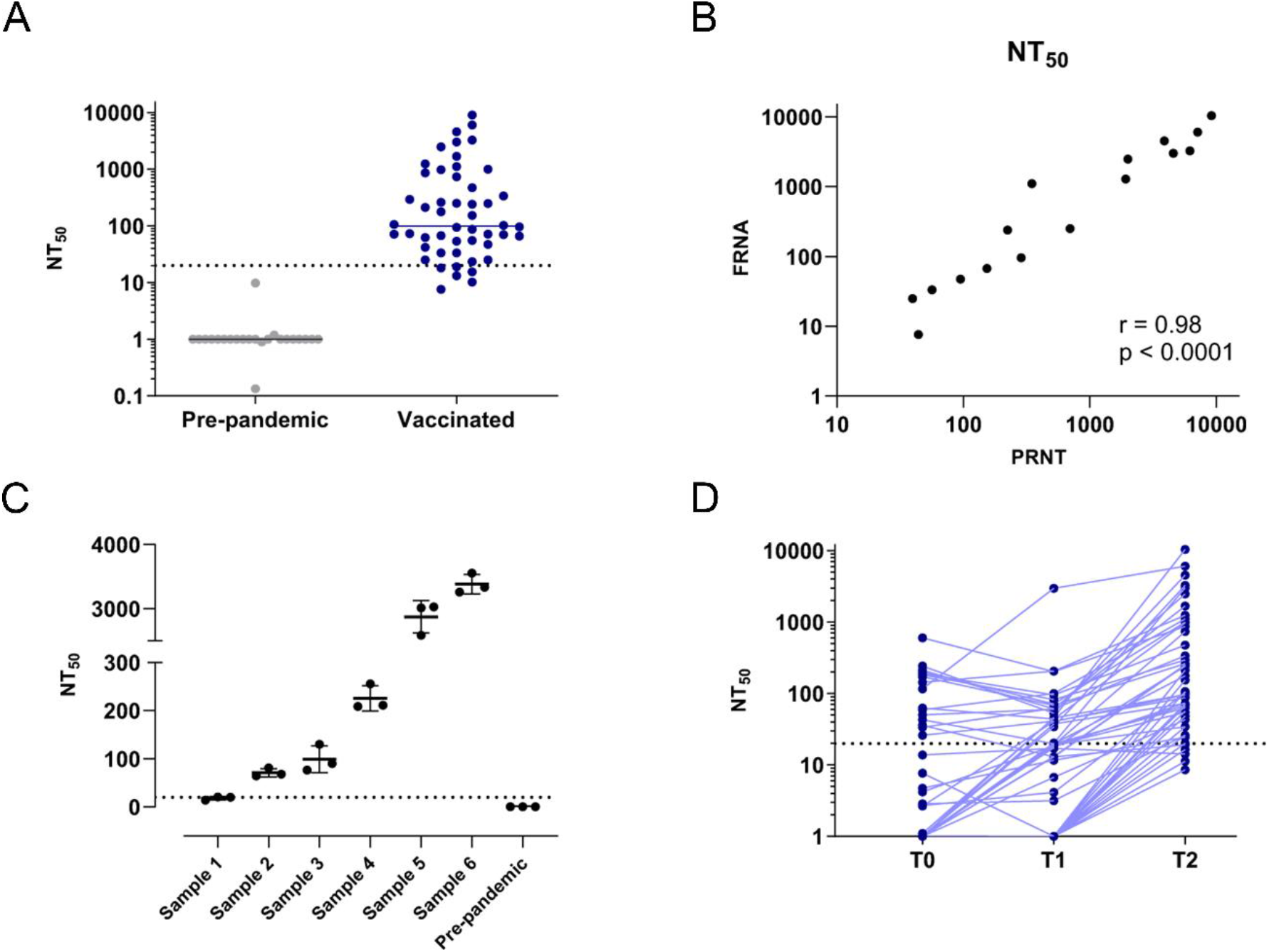
FRNA validation. A) Serum samples (a total of 74 samples; 24 pre-pandemic and 50 after the second dose vaccination) analysed by FRNA. B) Correlation between the PRNT and FRNA assays. A correlation of r = 0.98 was obtained (p < 0.0001). C) Reproducibility test. Six positive samples and one pre-pandemic were evaluated in three independent experiments, and NT50 was calculated. D) Paired samples from T0 (before vaccination), T1 (after the first dose), and T2 (after the second dose). Dashed lines represent the test cut-off: negative = NT50 < 20 (detection limit).

Since PRNT is the gold standard method for detecting nAbs, we tested a panel of positive samples using both techniques to compare antibody titers. Sixteen samples were selected from the panel to include a wide range of neutralization titers and samples from individuals who received different vaccines. Both techniques yielded similar antibody titers, with a correlation coefficient of r=0.98 (Figure 3B), showing that the newly developed FRNA provides reliable results within 48 hours considering experimentation and analysis time.

In order to evaluate the reproducibility of the FRNA, we analysed seven samples in three independent experiments. We included samples with high, intermediate, and low NT50 values and one pre-pandemic sample in the assays and observed low variability, which supports FRNA as a suitable methodology for evaluating anti-SARS-CoV-2 nAbs (Figure 3C).

Following the validation process, we analyzed pre-vaccination samples (T0) and samples from after the first and second vaccine doses (i.e., T1 and T2). We observed that 31% of samples were positive before vaccination, indicating prior SARS-CoV-2 infection. After the first dose, 39% of samples were positive for nAbs, increasing to 88% after two doses (Figure 3D). These results demonstrated that FRNA is a useful method for monitoring the kinetics of nAbs in the population, which can help support decision-making by health authorities regarding vaccine boosters and/or vaccine development.

### Production of SARS-CoV-2 mAbs and its use in fluorescent-based neutralization assay (FRNA)

Since the FRNA is reliant on immunofluorescence staining to determine NT50 titers, SARS-CoV-2 specific mAbs represent an important reagent for the success of the assay. In order to produce hybridomas that secrete anti-SARS-CoV-2 antibodies, Balb/C mice were immunized with the purified virus, and their splenocytes were fused with myeloma cells. Hybridoma screening and stabilization by freeze-thaw cycles resulted in 24 hybridomas positive for antibody secretion against SARS-CoV-2 (Sup. Figure 1). Based on cell growth, antibody secretion, and IFA staining patterns, three hybridomas were subjected to limiting dilution to obtain antibodies of monoclonal origin: 1F7, 17E12, and 25C3.

A characterization of these mAbs showed that they target different SARS-CoV-2 variants: 17E12 is reactive against the Wuhan, delta, and omicron variants, while 25C3 and 1F7 reacted against Wuhan, delta, omicron, and both gamma isolates (Figure 4A). mAb isotyping showed that 1F7 and 17E12 are IgG1 isotypes while 25C3 is an IgG2 isotype, all with *kappa* light chains.

**Figure 4.**
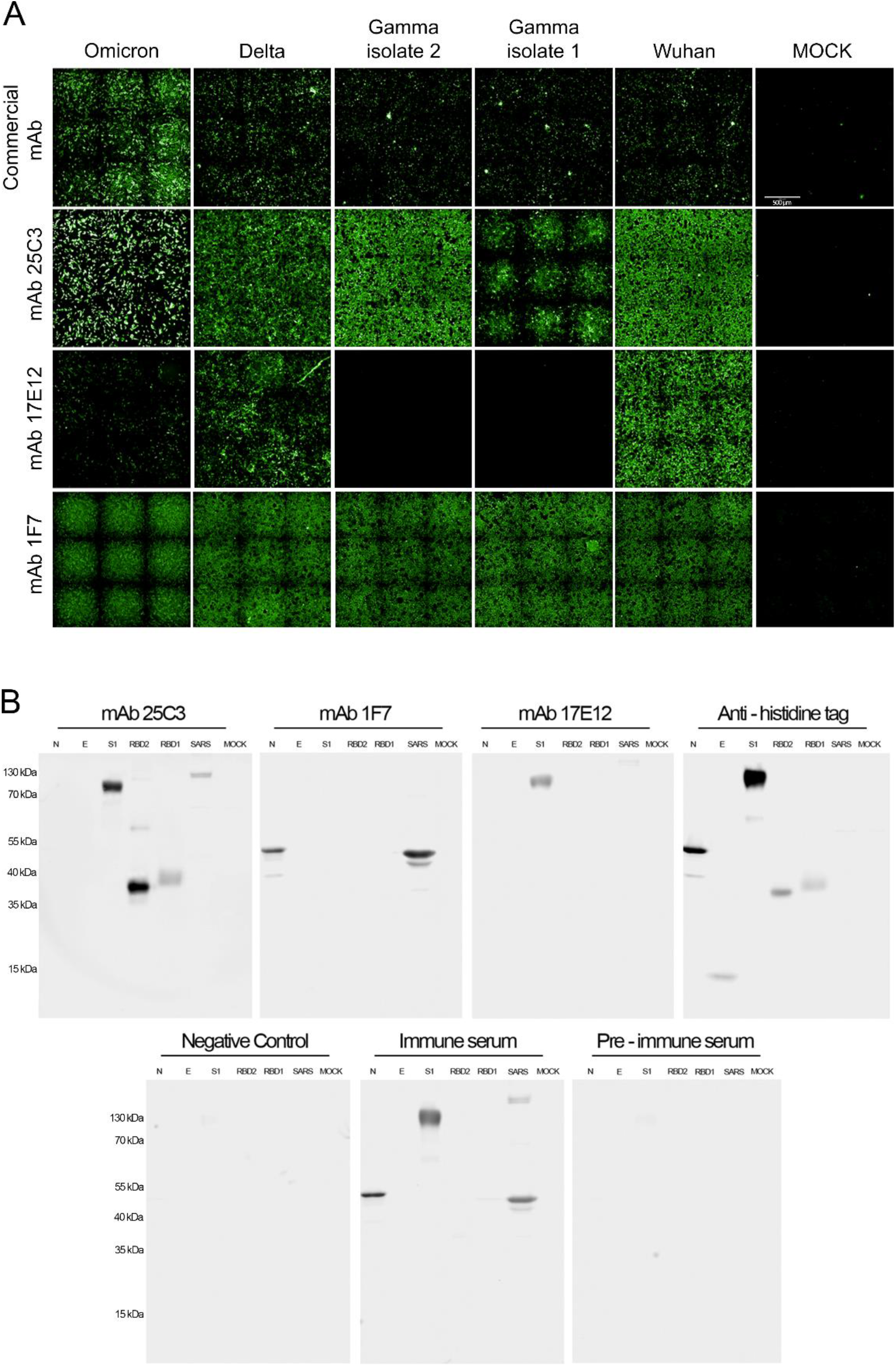
Characterization of anti-SARS-CoV-2 monoclonal antibodies. A) Reactivity by IFA of selected in-house mAbs (1F7, 25C3, and 17E12) against SARS-CoV-2 variants of concern in Vero E6 infected cells. B) Western blot analyses and characterization of selected mAbs against different SARS-CoV-2 recombinant proteins. Pre-immune serum was used as a negative control, post-immune serum was used as a positive control, and a non-related primary antibody was used as the negative control.

The reactivity of the mAbs to SARS-CoV-2 proteins was evaluated by WB using a lysate of Vero E6 infected cells and recombinant viral proteins. mAb 1F7 presents specificity to the N protein, as shown by the recognition of the rN protein and the ∼55 kDA band in the infected cell lysate. mAb 25C3 showed reactivity against the receptor binding domain (RBD) of both SARS-CoV and SARS-CoV-2 S1 protein. mAb 17E12 also showed reactivity against S1 but did not react against the RBD, indicating specificity to a different epitope (Figure 4B).

Based on the characterization of the in-house mAbs, 25C3 was selected to evaluate its applicability as a primary antibody in the FRNA. We submitted 24 samples from the panel of vaccinated individuals to the FRNA, as previously described. Aiming to determine if FRNA results with mAb 25C3 correlated with the gold standard method, PRNT, we compared NT50 titers obtained using both techniques and found a correlation of r=0.90 (Figure 5A). In addition, NT50 titers obtained from the FRNA using mAb 25C3 presented a strong positive correlation with the results obtained using the commercial mAb, with a coefficient of r=0.89 (Figure 5B). These results demonstrate that the in-house mAb 25C3 is a suitable reagent for the FRNA since its performance is comparable to the commercial mAb and results in similar NT50 titers to the PRNT.

**Figure 5.**
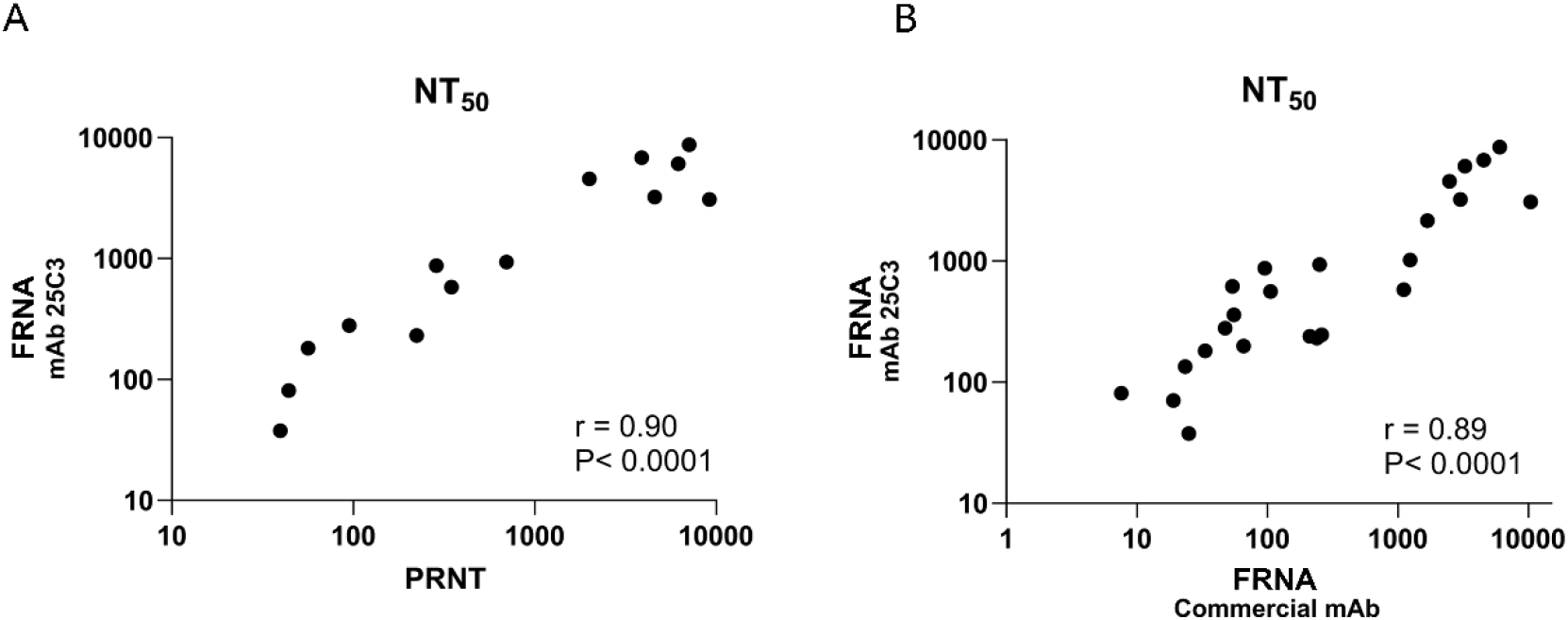
FRNA validation using in-house mAb 25C3. A) Correlation between standard gold method PRNT and FRNA titers using mAb 25C3. A correlation of r = 0.90 was obtained (p < 0.0001). B) Correlation between FRNA using commercial mAb and mAb 25C3. A correlation of r = 0.89 was obtained (p < 0.0001).

### In situ detection of SARS-CoV-2 by Immunohistochemistry

Since IHC is an important and widely used assay in medical research and clinical diagnostics, we validated one of the mAbs we developed in pulmonary tissue samples from COVID-19 deceased patients. We selected the 1F7 mAb, which targets the viral N protein, due to the high expression level during the infection cycle and its degree of amino acid identity among SARS-CoV-2 VOCs^15,16^. The results of all sample tests were positive when both the anti-SARS-CoV-2 commercial mAb (anti-Spike) and the 1F7 mAb were used. As depicted in Figure 6, both mAbs displayed specific immunostaining with minimal background noise. However, the reaction of the 1F7 mAb was found to be even more specific and without any nuclear background, as it was only present in the cytoplasm of pneumocytes and macrophages (Figure 6).

**Figure 6.**
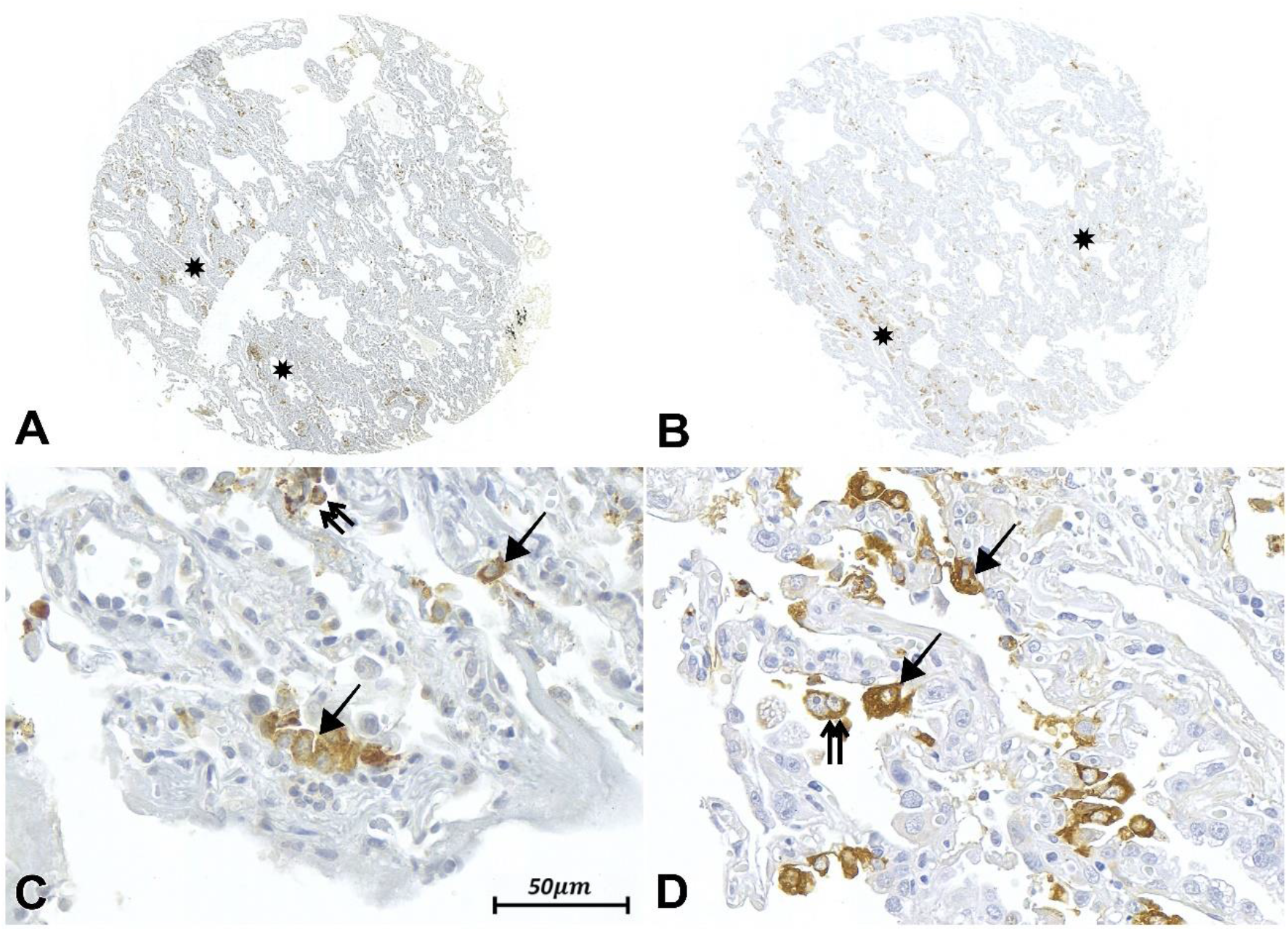
Immunohistochemistry assay using monoclonal antibodies to detect SARS-CoV-2 expression in the alveolar parenchyma of lung samples from COVID-19 patients. Panoramic photomicrographs in A and B demonstrate scattered areas in brown (*), which indicate positive immunoreactivity for the virus using Abcam and 1F7 mAbs, respectively. The Abcam commercial mAb (A) shows slight background staining. In C, a 200x magnification of the tissue expression with Abcam mAb reveals some overlapping of the stain with the nuclei of type II pneumocytes (black arrow) and alveolar macrophages (double arrows). However, in D, a 200x magnification of the tissue expression using 1F7 mAb (from the same sample area as C) clearly shows a granular and brown pigment in the cytoplasm of type II pneumocytes and alveolar macrophages without any nucleus overlap.

## Discussion

Since the COVID-19 pandemic began in late 2019, worldwide efforts have been made by the scientific community to assist health authorities with their decision-making. In addition to fast differential diagnosis and vaccine development, there was an urgent need for scalable and high-throughput methods to evaluate the immunological status of infected and/or vaccinated people. Assays that could serve as protection indicators are important to guide public health decisions, including the development of new vaccines or adaptations of current vaccination strategies.

Humoral immunity is essential to control viral infection. In the case of SARS-CoV-2, it was recently demonstrated that most infected individuals seroconverted (IgG and IgM production) within 19 days after symptom onset^17^. Despite the correlation between nAbs levels and protection against secondary infections and severe disease^18^, it was suggested by predictive models that humoral immunity may wane with time as a result of declining nAb levels^19^. Recent studies demonstrated a significant decrease of nAbs in sera six months after infection and eight months after mRNA vaccination^20,21^. In addition to waning immunity, the emergence of SARS-CoV-2 variants with the potential to escape vaccine-derived immune protection has caused concern. It has been observed that the omicron variant escapes from immunity within as little as four weeks after the second dose of mRNA vaccination^22^. Omicron subvariants that have recently emerged appear to escape from neutralization after monovalent and/or bivalent vaccine boosters^23,24^.

PRNT is well established as the gold standard method for quantifying neutralization titers against viral infections^25^, but it has several disadvantages. The assay has a turnaround time of one week and is usually carried out in 24-well plates, which requires large sample volumes and limits the number of samples that can be analyzed per plate. Moreover, PRNT is a laborious technique, requiring trained personnel to manually quantify plaque-forming units. These limitations prevent its use to assess neutralization titers on a large scale, which is often needed during epidemics when there is a demand for serological surveys. To overcome these limitations, we developed a potential scalable and high-throughput fluorescent-based neutralization assay to evaluate SARS-CoV-2 nAbs. Because the assay relies on immunostaining, we also produced in-house anti-SARS-CoV-2 mAbs and demonstrated its applicability in the FRNA and IHC assays.

Our assay combines classical neutralization principles with an automated method using a high-content imaging system. From seeding cells up to titer calculation, results are obtained in less than 48 hours, representing time savings of around 70% compared to the gold standard PRNT. Its 96-well format enables the automation of assay steps and allows the processing of at least six samples/plate simultaneously compared to one sample/plate in the classical PRNT. Moreover, the FRNA determines antibody titers based on infection percentages from IFA images, which are calculated automatically. Therefore, it eliminates the subjectivity of PRNT quantification due to the manual counting of viral plaques. Our validation results demonstrated that the FRNA is highly applicable to detect nAbs post-vaccination and presented a high correlation with the PRNT (r = 0.98) and low inter-assay variability. In addition, the assay proposed here uses the live virus, which increases accuracy and makes it readily adaptable to quantify nAbs against emerging SARS-CoV-2 VOCs. On the other hand, the assay can only be performed in a biosafety level 3 (BSL3) facility and relies on high-content imaging equipment, which is costly and may not be easily accessed by many laboratories. Alternatively, the images can be obtained in a convectional fluorescence microscope and analyzed by independent software.

In an attempt to overcome PRNT limitations, adaptations were recently developed using SARS-CoV-2 pseudoviruses. These genetically modified viruses express the SARS-CoV-2 spike protein on the core skeleton of a non-coronavirus system such as vesicular stomatitis virus (VSV), while also encoding a quantifiable reporter gene^26^. This technique quantifies viral foci by luminescence of the reporter gene in plate readers, increasing throughput. Another advantage of this method is that it can be performed in a biosafety level-2 laboratory (BSL-2). However, the production and validation of pseudoviruses are complex, requiring highly skilled operators, and assay readout may need specialized equipment. Because pseudoviruses are typically nonreplicating or single-cycle viruses that only express the spike protein from SARS-CoV-2, spike density on the virion surface and replication kinetics differ from the live virus^27,28^. Therefore, correlation with live virus assays and assay sensitivity may vary greatly between different pseudovirus systems. For instance, it has been observed that some HIV-1 and VSV-based pseudovirus assays were less sensitive than wild-type neutralization assays, especially when evaluating weakly neutralizing plasma^29^.

MAbs represent a crucial but costly reagent for the success of the FRNA and are widely used in other laboratory techniques such as flow cytometry, WB, ELISA, and IHC. In addition to high cost, the increased demand for reagents during the SARS-CoV-2 pandemic caused global shortages of several products, including mAbs used for research, diagnostics, and treatment purposes^30,31^. To overcome this issue and due to our previous experience in mAb development^13^, we developed a panel of anti-SARS-CoV-2 mAbs. Using hybridoma technology, we generated stable cell lines that secrete antibodies and therefore provide an unlimited supply of anti-SARS-CoV-2 mAbs.

The mAbs we developed target different SARS-CoV-2 proteins (RBD, S1, and N proteins), which is advantageous for adapting the assay to other variants. The mAb 25C3 binds to RBD, which has been shown to strongly interact with the angiotensin-converting enzyme 2 (ACE2) of humans and bat cells^32^. Although mutations could impair the binding of antibodies to RBD^33^, we demonstrated that mAb 25C3 was able to target all SARS-CoV-2 variants tested. It is important to highlight that new variants could eventually escape mAb 25C3 binding; therefore, we identified other mAbs that could be used as alternatives in the FRNA. In the panel we tested, mAbs 1F7 and 17E12 were able to target the N and S1 proteins, respectively. The N protein is highly expressed during infection^15^, which makes the mAb 1F7 a candidate to be used in diagnosis, as demonstrated by IHC assays in SARS-CoV-2 positive pulmonary tissue samples. When used in parallel with a commercial SARS-CoV-2 mAb, mAb 1F7 was more specific while maintaining minimal background noise. These results demonstrate the applicability of mAb 1F7 in IHC, an important complementary diagnostic assay when other samples are unavailable or not collected within an adequate timeframe.

In conclusion, the fluorescence reduction neutralization test we developed represents a significant advance compared to the gold standard PRNT. The assay had a high correlation with PRNT, with a 70% reduction in turnaround time and an automated system that allows high-throughput sample processing while eliminating the subjectivity of PRNT. We also produced anti-SARS-CoV-2 mAbs that target different viral proteins and recognize VOCs, demonstrating their applicability in the neutralization and IHC assays. The development of these mAbs reduced assay costs, guaranteeing the independence of the importation bureaucracy system and ensuring their availability in unlimited quantities.

## Acknowledgments

The authors thank Wagner Nagib de Souza (ICC/Fiocruz-PR) for providing the infographic in Figure 1A. Thanks also to Dr. José Luiz Proença Módena for gently providing the Omicron variant.

## Figures

**Figure S1.**
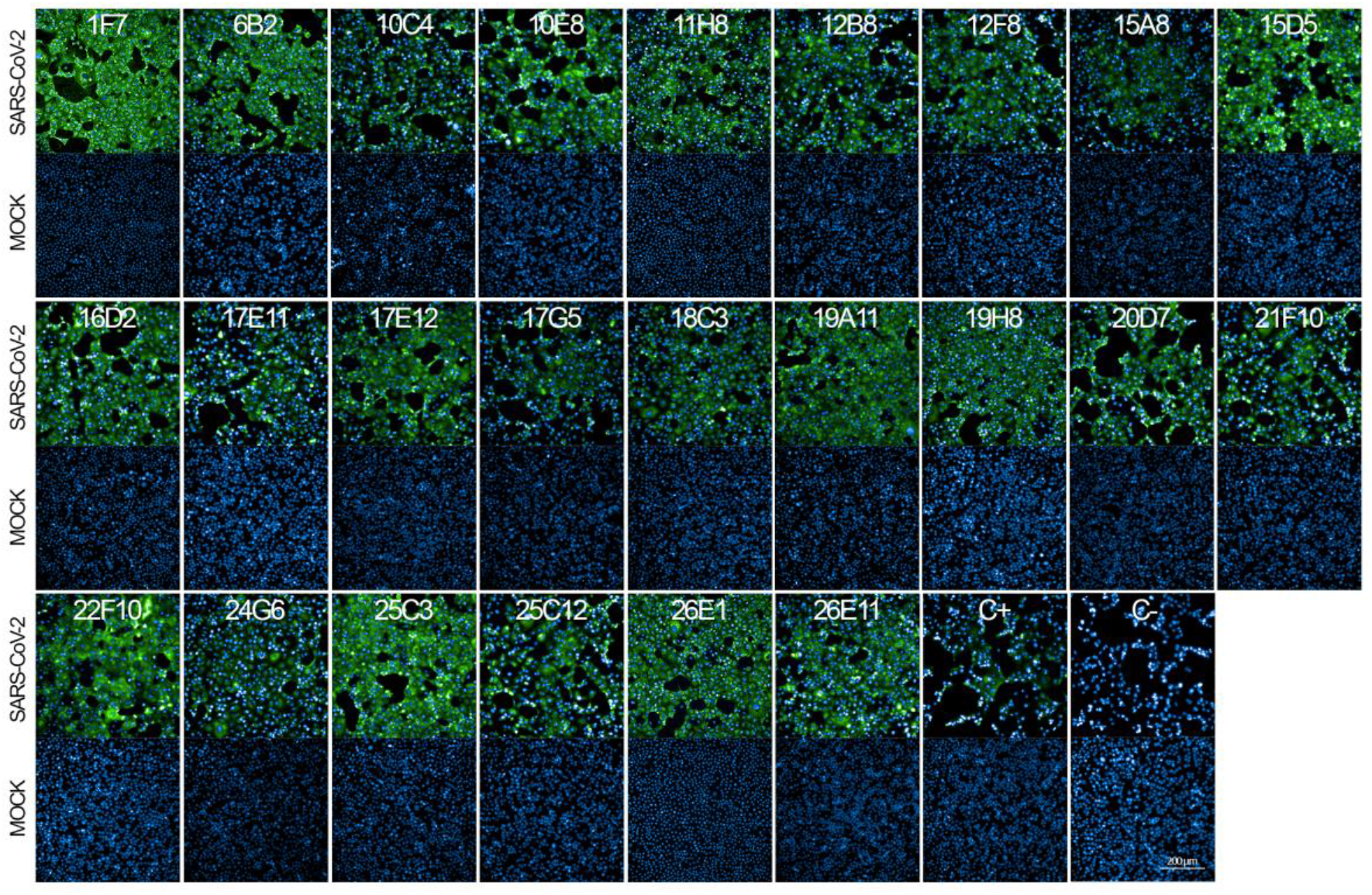
Indirect immunofluorescence assay of polyclonal hybridomas secreting antibodies against SARS-CoV-2. Vero E6 cells were infected with SARS-CoV-2, fixed, and incubated with hybridoma supernatant, followed by an anti-mouse IgG antibody conjugated with Alexa-fluor 488. An anti-spike commercial Ab were used as positive and negative controls, respectively. Antibodies were also tested against MOCK (uninfected) cells. Images were obtained with the CLS High-Content Imaging System (Perkin Elmer).

